# normR: Regime enrichment calling for ChIP-seq data

**DOI:** 10.1101/082263

**Authors:** Johannes Helmuth, Na Li, Laura Arrigoni, Kathrin Gianmoena, Cristina Cadenas, Gilles Gasparoni, Anupam Sinha, Philip Rosenstiel, Jörn Walter, Jan G. Hengstler, Thomas Manke, Ho-Ryun Chung

**Author notes:** Corresponding author: Ho-Ryun Chung, Max Planck Institute for Molecular Genetics, Ihnestrasse 63–73, 14195 Berlin, Germany.

## Abstract

ChIP-seq probes genome-wide localization of DNA-associated proteins. To mitigate technical biases ChIP-seq read densities are normalized to read densities obtained by a control. Our statistical framework “normR” achieves a sensitive normalization by accounting for the effect of putative protein-bound regions on the overall read statistics. Here, we demonstrate normR’s suitability in three studies: (i) calling enrichment for high (H3K4me3) and low (H3K36me3) signal-to-ratio data; (ii) identifying two previously undescribed H3K27me3 and H3K9me3 heterochromatic regimes of broad and peak enrichment; and (iii) calling differential H3K4me3 or H3K27me3-enrichment between HepG2 hepatocarcinoma cells and primary human Hepatocytes. normR is readily available on http://bioconductor.org/packages/normr

## Introduction

Chromatin Immunoprecipitation followed by high-throughput sequencing (ChIP-seq; Johnson et al. 2007) is a widely used method for the genome-wide localization of DNA-associated proteins, such as transcription factors or histone modifications. In brief, after crosslinking with formaldehyde the chromatin is sheared and the resulting chromatin fragments are enriched by immunoprecipitation for the protein of interest. The precipitate is reverse-crosslinked to obtain DNA fragments, which are amplified and then sequenced. The reads generated in this way are then aligned to a reference genome and genomic loci bound by the protein are inferred by an accumulation of sequencing reads. Due to the genome-wide scalability and cost-efficiency of ChIP-seq, hundreds of distinct proteins and their modifications have been assayed to study underlying mechanisms of molecular function in different cell types (ENCODE Project Consortium 2012,Roadmap Epigenomics Consortium et al. 2015). Consequently, a huge resource of protein location information is available to be readily integrated into studies at hand.

ChIP-seq data are used to characterize transcription factor binding sites Thomas-Chollier et al. 2012, chromatin landscapes (Mammana and Chung 2015; Perner et al. 2014) or functional elements, like enhancers (Heintzman et al. 2007; 2009). Specifically, most ChIP-seq experiments aim to study protein binding sites in the context of gene regulation. For example, the lineage-specific binding of transcription factors orchestrates differentiation pathways (Tsankov et al. 2015). Furthermore, ChIP-seq signals of histone modifications are predictive for promoter activity (Karlić et al. 2010) and enhancer competence (Bonn et al. 2012).

The identification of regions bound by a protein of interest requires the discrimination of enrichment against background. Intuitively, a high number of ChIP sequencing reads should map to protein-bound regions, where the average number of reads in these regions depends on the “binding mode” of the protein of interest. For example, transcription factors and certain histone modifications, such as H3K4me3, are characterized by a localized read accumulation with a high signal-to-noise ratio (Sims et al. 2014). Some histone modifications, such as H3K9me3, H3K27me3, and H3K36me3 are characterized by a more delocalized read accumulation with a substantially lower signal-to-noise ratio.

Technical biases introduced during the ChIP-seq procedure lead to accumulation of reads in regions that are devoid of the protein (Vega et al. 2009; Jain et al. 2015; Meyer and Liu 2014). These biases arise by copy number variations, sequencing biases, mapping ambiguities, and the chromatin structure (Vega et al. 2009; Flensburg et al. 2014). These biases are also discernable in control experiments, *i.e.* they can be accounted for by comparing the ChIP read coverage to a control experiment without specific enrichment, such as the input chromatin to the ChIP.

The comparison of the read counts in the ChIP to those in the control requires normalization to account for, both, the differences in the sequencing depth, and the effects of enrichment by the ChIP. Ideally, such a normalization should yield a normalization factor that corrects the average ratio between ChIP- and control read counts in background regions (Liang and Keles 2012; Xu et al. 2010; Diaz et al. 2012). Thus, a proper normalization requires the identity of background regions. On the other hand, the discrimination of enriched and background regions requires normalization itself – normalization and discrimination of enrichment against background are two faces of the same coin.

Earlier approaches estimate the normalization factor either by the ratio of sequencing depths (*e.g.* MACS (Zhang et al. 2008) and DFilter (Kumar et al. 2013)), by the ratio of ChIP- and control read counts summed over *ad hoc* chosen background regions with fixed width (e.g. CisGenome (Ji et al. 2008), SPP (Kharchenko et al. 2008) and MUSIC (Harmanci et al. 2014)), or by identifying background regions and their width using a data-driven approach (e.g. NCIS (Liang and Keles 2012) or SES (Diaz et al. 2012)). After normalization these approaches identify enriched regions and equate them to protein binding sites or modifications. All these approaches discriminate a single signal regime from the background. However, a qualitative separation of this signal regime, *e.g.* into moderately and highly enriched regimes, could distinguish genomic loci that are bound by the protein in only a subpopulation of cells in the sample from those that are bound in the majority of cells in the sample. Those analyses of ChIP-seq sample heterogeneity cannot be performed using existing methods.

The discrimination of signal against background is not only required to determine protein binding sites it is also required for identifying regions that are differentially bound in two conditions, *e.g.* control and disease. Most approaches (*e.g.* MACS (Zhang et al. 2008) aimed at identifying differentially bound regions concentrate on the modeling of condition-specific exclusive enrichment. In addition, other methods (Xu et al. 2008; Allhoff et al. 2014; Heinig et al. 2015) employ a three-state Hidden Markov Model to additionally identify condition-specific changes of signal within regions of concurrent ChIP enrichment. Therein, a computationally intensive training is done to learn a hidden state representation of the data. Consequently, the regional ChIP read coverage is “interpolated”/”smoothed” based on the read coverage in adjacent genomic loci. This data abstraction sacrifices a statistically sound null hypothesis.

Here, we describe a data-driven robust and broadly applicable approach for simultaneous normalization and difference calling in ChIP-seq data called normR (recursive acronym: “normR obeys regime mixture rules”). normR models ChIP- and control read counts by a binomial mixture model. One component models the background, while one or more other components model the signal. As a proof of principle, normR is applied in three scenarios: Firstly, we show that normR achieves robust enrichment calling for both high (H3K4me3) and low (H3K36me3) signal-to-noise ratio ChIP-seq data. High specificity and sensitivity of normR is confirmed by functional outputs like gene expression and DNA methylation state. Secondly, we use normR to characterize two previously undetectable enrichment regimes for H3K27me3 and H3K9me3 in hepatocarcinoma HepG2 cells. Finally, the translational normR approach is shown to confidently call differences between primary human hepatocytes and HepG2 cells for both high (H3K4me3) and low (H3K27me3) signal-to-noise ratio histone modification ChIP-seq data. Here, we uncover potential epigenetic alterations introduced by the cancer-associated immortalization of primary liver cells. Thus, normR is a versatile tool that can identify enriched regions, distinct enrichment regimes and differences between conditions using a simple binomial mixture model and robust statistics.

## Results

### The normR Framework

During a ChIP experiment antibodies are used to enrich chromatin fragments carrying the protein of interest from a population of fragments obtained by sonication of chromatin. These antibodies bind preferentially but not exclusively to protein-DNA complexes. Hence, ChIP only enriches rather than selects protein containing chromatin fragments. Bearing this in mind, ChIP can be envisioned as a sampling process where the probability to draw a fragment depends on the presence or absence of the protein. If present, the probability is high, if absent, the probability is lower but not zero. The spatial distribution of the fragments sampled in this way is then estimated by mapping the sequenced ends (reads) of these fragments to the reference genome.

To infer regions bound by a protein of interest the read densities obtained by ChIP-seq experiment are compared to the corresponding counts obtained by a control experiment *e.g.* by sequencing the sonicated chromatin (input). A region should be called “enriched by the ChIP” only if the number of reads from the ChIP is sufficiently greater than that expected relative to the control. Such an approach addresses a number of systematic biases, like copy number variations, sequencing biases, mapping ambiguities and chromatin structure (Vega et al. 2009; Meyer and Liu 2014; Flensburg et al. 2014). To this end, a proper normalization of the read count densities is essential: For example, if we sequence twice as many reads in the ChIP than in the control, the read counts per region in the ChIP should be greater than in the control. In the absence of enrichment by the ChIP, we expect twice as many reads per region in the ChIP than in the control. In the presence of enrichment by the ChIP, the read counts in the region associated with the protein should be much higher than in the control, but what happens to the read counts in the remaining regions?

Sequencing the ChIP and control libraries is a multinomial sampling process, which induces dependencies between the regions. As the total number of reads obtained from one sequencing run is fixed and finite, the increase of reads in some regions due to ChIP enrichment leads to a decrease in remaining regions, *i.e.* background regions *B.* Returning to our example, this implies that the number of reads in non-enriched regions in the ChIP should be less than twice the number from the control. In particular, the normalization factor *C*_*B*_ is less than two which relates the number of reads in ChIP-seq *s*_*i*_ to the ones in control *r*_*i*_ by 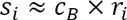 for background regions *i ∈ B*. *c*_*B*_ depends on the average enrichment achieved by the ChIP and the number of enriched regions — it shrinks as, both, the number of enriched regions and the level of enrichment in these regions increases. Critically, *C*_*B*_ is required to define a statistically sound Null hypothesis for testing whether the observed ChIP read counts are sufficiently greater than expected given the control. Moreover, the more regions are enriched, the lower the signal-to-noise ratio becomes at a fixed sequencing depth (Sims et al. 2014). The estimation of *C*_*B*_ requires the identity of background regions, albeit the identification of the background requires normalization itself. Thus, ChIP-seq normalization and the identification of enriched regions are two sides of the same problem.

To tackle this problem we model the read counts from the ChIP and control by a binomial mixture model (Methods; Figure 1). In its simplest incarnation we use two components, *i.e.* background and enriched, to normalize and call enrichment over the control (referred to as “enrichR”). The model has in total three free parameters, *i.e. θ*_*B*_, *θ*_*E*_ and *π*_*B*_ · *θ*_*B*_ and *θ*_*E*_ represent the expected fraction of reads in the ChIP over the sum of reads from ChIP and control per region for the background and the enriched regions, respectively. *π*_*B*_ is the proportion of regions that belong to the background *π*_*B*_ (the proportion of regions that are enriched is simply *π*_*E*_ *=* 1 – *π*_*B*_). Given this model we derive the following likelihood function:

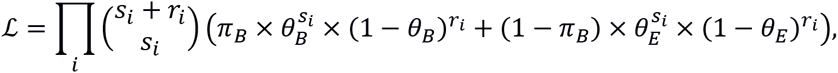

where *s*_*i*_ (*r*_*i*_) corresponds to the number of reads in the ChIP (control) for regions *i =* 1, …,*n*. We fit these parameters using the expectation-maximization algorithm (Dempster et al. 1977) on the closed form solution (Methods). From the discussion above we expect that 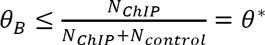, where *N*_*chIP*_ (*N*_*control*_) is the total number of reads in the ChIP (control) and *θ\** denotes the expected fraction of reads from ChIP-seq taking into account only sequencing depth. Equality holds only in case of no enrichment, or *π*_*B*_ = 1. The last implicit “parameter” is the definition of regions. We use non-overlapping fixed width regions because it is robust and appropriate for most downstream analyses (Ernst and Kellis 2012; Nair et al. 2014; Ramírez et al. 2014; Mammana and Chung 2015).

**Figure 1.**
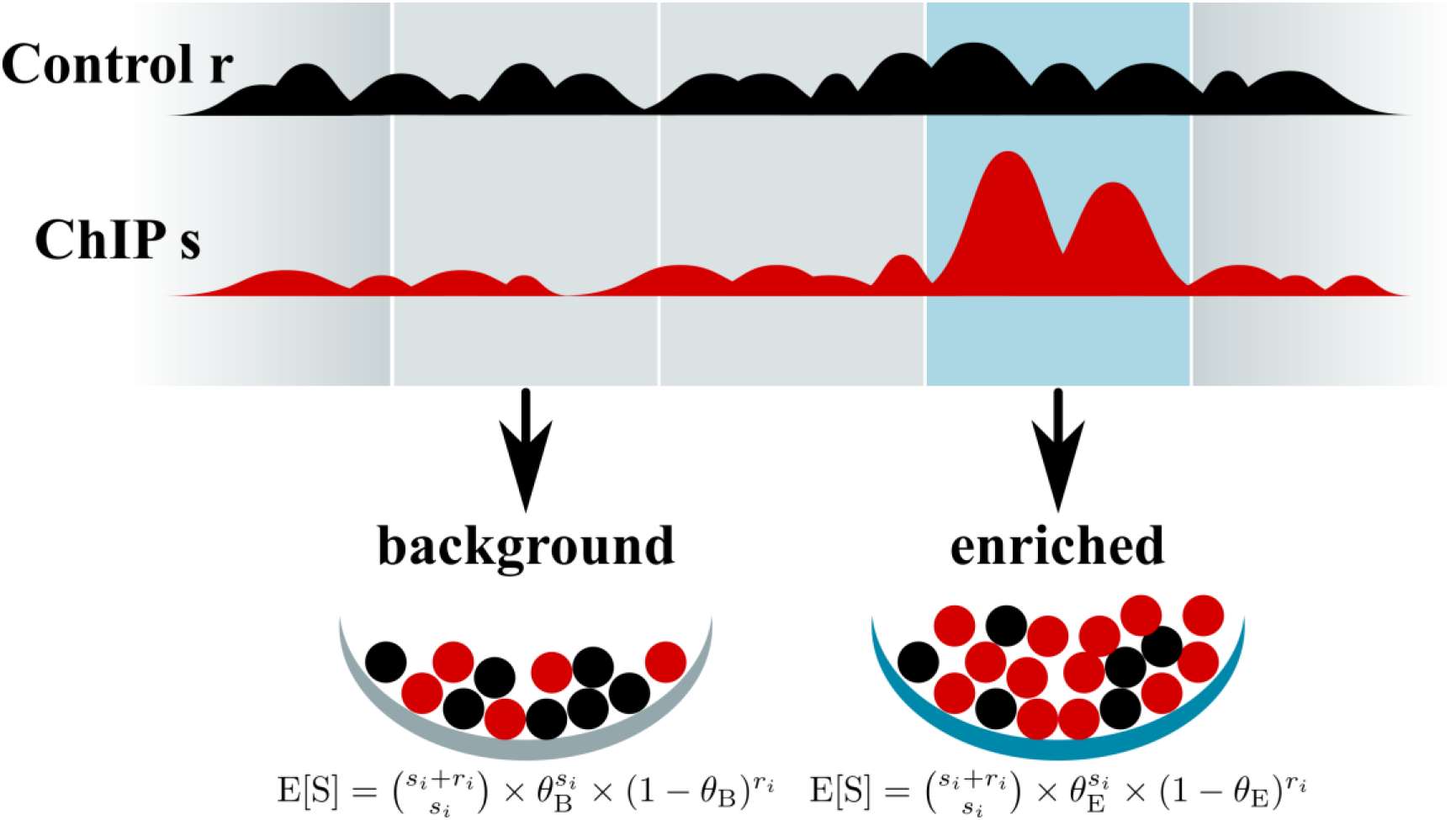
The normR Framework. Reads in control *r* and ChIP *s* are modeled as a binomial mixture model with multiple components. Here, two components model the expected fraction of reads in the ChIP over the sum of reads from ChIP and control per region for background *θ*_*B*_ and the enriched *θ*_*E*_. By accounting for the effect of ChIP enrichment on the background read statistics a statistical sound Null hypothesis is formed.

The identification of enriched regions across the genome is based on the fitted model: Given the control read count, the ChIP read count in each region is compared to the expected read count under the fitted background model. Using a binomial test statistically significant deviations from the background model are recovered. The null distribution of p-values from a binomial test is discrete and impedes the correction for multiple testing. By filtering out low power tests (*i.e.* low count regions) with the T method (Dialsingh et al. 2015), the p-value distribution becomes more uniform and the p-values can be adjusted for multiple testing. Filtered p-values are then transformed to q-values (Storey 2002). Enriched regions are reported if they fall below a user-specified threshold.

In addition to enrichR, we provide two augmented realizations of normR (Methods): (i) “regimeR” models multiple enrichment components defined by 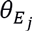 with *j* = 1,…, *m* to identify ChIP enrichment regimes; and (ii) “diffR” models the expected fraction of reads in a depleted (control-enriched) component defined by *θ*_*D*_ in addition to *θ*_*B*_ and *θ*_*E*_ yielding a direct comparison of two ChIP experiments. After assessing significance against *θ*_*B*_ every region is assigned to a component by *Maximum a posteriori* assignment.

Based on the fitted binomial mixture model the normalized ChIP signal *e*_*i*_ is calculated by dividing the read counts from ChIP-seq by those from the control scaled by the normR enrichment factor 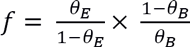. To account for noise in low power regions, we regularize *e*_*i*_ by adding pseudocounts to the number of ChIP-seq and Input-seq reads resulting in

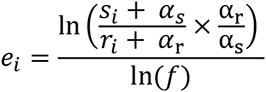

where 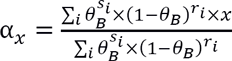 represents the average read count for *x* given the normR-fitted background model.

We have implemented normR in C++ and R (R Core Team 2015). normR is available on Bioconductor at http://bioconductor.org/packages/normr.

### Enrichment Calling in Low and High Signal-To-Noise Ratio Settings with enrichR in Primary Human Hepatocytes

To illustrate the enrichment calling based on a robust background estimation, we applied enrichR to two ChIP-seq experiments against H3K4me3 and H3K36me3 in primary human hepatocytes. H3K4me3 correlates with promoter activity and DNA-hypomethylation (Ooi et al. 2007; Long et al. 2013; Hu et al. 2009) and exhibits a high signal-to-noise ratio (Supplemental Fig. 1). H3K36me3 represents a lower signal-to-noise characteristics (Supplemental Fig. 1) and is associated to transcriptional elongation in the body of transcribed genes (Kim et al. 2007) as well as DNA-hypermethylation (Baubec et al. 2015). We performed enrichR analyses on the ChIP-seq data against Input-seq (Methods). The enrichment calls by enrichR were compared to peaks called by six popular peak calling tools ChIP-seq data: MACS2 (Feng et al. 2012), DFilter (Kumar et al. 2013), CisGenome (Ji et al. 2008), SPP (Kharchenko et al. 2008), BCP (Xing et al. 2012) and MUSIC (Harmanci et al. 2014).

As a first assessment, we inspected the coverage and enrichment/peak calls for H3K4me3 and H3K36me3 ChIP-seq in the vicinity of the Glucose-6-Phosphate Isomerase gene (GPI, Figure 2A) — a housekeeping gene that is highly expressed in all cell types (Eisenberg and Levanon 2013). GPI was also expressed in primary human hepatocytes as measured by RNA-seq and showed a characteristic chromatin signature of transcription, *i.e.* H3K4me3 and H3K36me3 in the promoter and the gene body, respectively. All tested methods identified these characteristic enrichments at the GPI locus. Moreover, the promoter of the WTIP gene was detected as H3K4me3-enriched by all methods. Together with the measured shallow coverage of RNA-seq reads along its gene body this indicated that WTIP is expressed suggesting a genuine H3K36me3 enrichment in its gene body. Interestingly, this minute H3K36me3 enrichment was exclusively recovered by enrichR.

**Figure 2.**
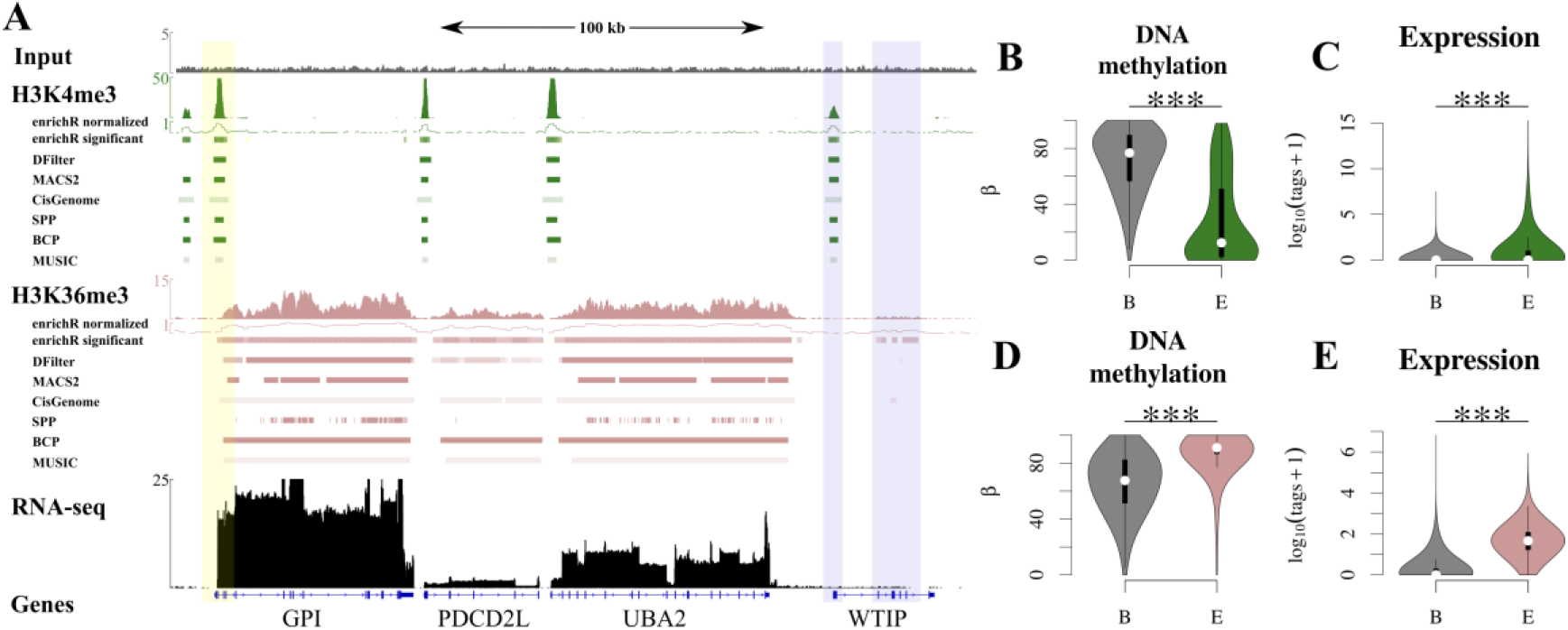
Enrichment Calling in Low and High Signal-To-Noise Ratio Settings with enrichR in Primary Human Hepatocytes. (A) Input (grey), H3K4me3 (green, high signal-to-noise ratio), H3K36me3 (rose, low signal-to-noise ratio) and RNA-seq (black) barplots indicate coverage proximal to the human Glucose-6-Phosphate Isomerase (GPI, yellow overlay) locus on chromosome 19 in Primary Human Hepatocytes (PHH). Enrichment calls are indicated as colored boxes below respective tracks for enrichR, DFilter, MACS2, CisGenome’s SeqPeak and SPP. The WTIP gene (blue overlay) had detectable H3K4me3 enrichment at its promoter and minute H3K36me3 is recovered solely by enrichR. (B-C) enrichR H3K4me3-enriched regions were DNA-hypomethylated (B) and expressed as measured by CAGE (C). (D-E) enrichR H3K36me3-enriched regions were DNA-hypermethylated (D) and expressed as measured by RNA-seq (E).

Genome-wide enrichR called H3K4me3-enrichment in 142,451 500 base-pair (bp) regions in primary human hepatocytes, corresponding to 45,522 consecutive regions representing ~3% of the mappable genome (71.2Mb). The identified regions were characterized by low levels of DNA methylation (Figure 2B), in line with the idea that H3K4me3 represses DNA methylation (Ooi et al. 2007; Long et al. 2013; Hu et al. 2009). Furthermore, H3K4me3-enriched regions recovered by enrichR showed a higher density of CAGE-tags than the background (Figure 2C) indicating that they serve as active transcriptional start sites (TSSs) in this cell type. In fact, enrichR H3K4me3-enriched regions showed a statistically significant overlap with annotated TSSs (odds-ratio = 25.04, Fisher’s signed exact test, P ≤ 0.001, Supplemental Table 1). Together these observations support that enrichR identifies *bona fide* H3K4me3-enriched regions.

The comparison of enrichR enriched regions to MACS2, DFilter, CisGenome, SPP, BCP and MUSIC peaks revealed a substantial overlap at *FDR* = 0.1 indicating that for H3K4me3 in this dataset all six methods work well, although in terms of covered bp DFilter (39.8Mb) and CisGenome (38.7Mb) called almost two-fold fewer regions than the other tools (mean=65.3Mb; Supplemental Note, Supplemental Fig. 2A, Supplemental Table 2).

For H3K36me3 enrichR identified 559,560 1 kilo base-pair (kb) windows as enriched, corresponding to 85,293 consecutive regions representing ~20% of the mappable genome (599.6Mb). H3K36me3-enriched regions recovered by enrichR showed high levels of DNA methylation (Figure 2D), in line with the observation that H3K36me3 recruits DNMT3B leading to *de novo* DNA methylation (Baubec et al. 2015). Furthermore, these regions showed significantly higher RNA-seq read coverage than background regions (Wilcoxon-signed-rank test P ≤ 0.001, Figure 2E), in line with the idea that H3K36me3 covers the gene body of transcribed genes Kim et al. 2007. Furthermore, enrichR H3K36me3-enriched regions showed a statistically significant overlap with annotated transcripts (odds-ratio = 17.06, Fisher’s signed exact test, P ≤ 0.001, Supplemental Table 1). These results support that enrichR also identifies *bona fide* H3K36me3-enriched regions.

When compared to enrichR results, far less H3K36me3-enriched regions were reported by MACS2 (407.7Mb), BCP (396.5Mb), MUSIC (402.3Mb) and by especially DFilter (87.8Mb), SPP (25.1Mb) and CisGenome (36.4Mb), even when configured for detection in low signal-to-noise ratio settings (Methods). Almost all of these regions (MACS2: 399.1Mb; 97.9%, DFilter: 87.8Mb; 100%; CisGenome: 36.4Mb; 100%; SPP:24.2Mb; 96.7%; BCP:386.8Mb; 97.6%; MUSIC:382.6Mb; 95.1%) were recovered by enrichR which leads to very few exclusive regions for the benchmark methods (Supplemental Fig. 2B). Regions called exclusively by enrichR (93.6Mb; 16.7%) were characterized by a median distance of >2kb to peaks recovered by other methods (Supplemental Fig. 2C). Furthermore, these regions showed significantly higher DNA-methylation levels and transcriptional activity than background regions suggesting once more a genuine H3K36me3 enrichment (Wilcoxonsigned-rank test P ≤ 0.001, Supplemental Fig. 2D-E).

Next, we studied accuracy of H3K36me3-enrichment peak calls. Because there is no genome-wide ChIP-seq benchmark set on-hand, we defined a gold standard for each method based on a consensus vote among the six remaining tools (Kinkley et al. 2016) (Supplemental Note): At FDR 0.1 DFilter and CisGenome achieved both highest precision (1.00), while enrichR had the highest recall (0.997) and BCP had the highest F_2_-score (0.631; Supplemental Table 2). enrichR which called almost all regions of the five tools combined had a recall-weighted F_2_-score of 0.533 compensating its menial precision (0.186) at q-value ≤ 0.1 with a superior recall. In fact, enrichR has the highest precision at recall ≤ 0.9 indicating that the consensus vote defined gold standard does not contain many enrichR-exclusive regions at q-value ≤ 0.1 (Supplemental Fig. 3). In a second assessment, we studied the validity of tool-specific regions, *i.e.* the peak calls not represented in the gold standard. To this end we defined a unified gold standard of H3K36me3-enrichment, *i.e.* the union of seven tool-specific gold standards, and seven sets of tool-specific regions (Supplemental Note). For all methods, the unified gold standard exhibited a significantly higher enrichment (fold change over Input) than tool-specific regions for enrichR, MACS2, SPP, BCP and MUSIC (Wilcoxon-signed-rank test; P ≤ 0.01; Supplemental Fig. 2F). Among these, enrichR had the most tool-specific regions (205,064; 36.6%) and showed significantly higher enrichment as well as read coverage than background regions (Supplemental Fig. 2G). Furthermore, enrichR-specific regions were remote from unified gold standard regions (median=14Mb; Supplemental Figure 2H) and, yet, still overrepresented in annotated gene bodies (odds-ratio = 13; Supplemental Table 1).

**Figure 3.**
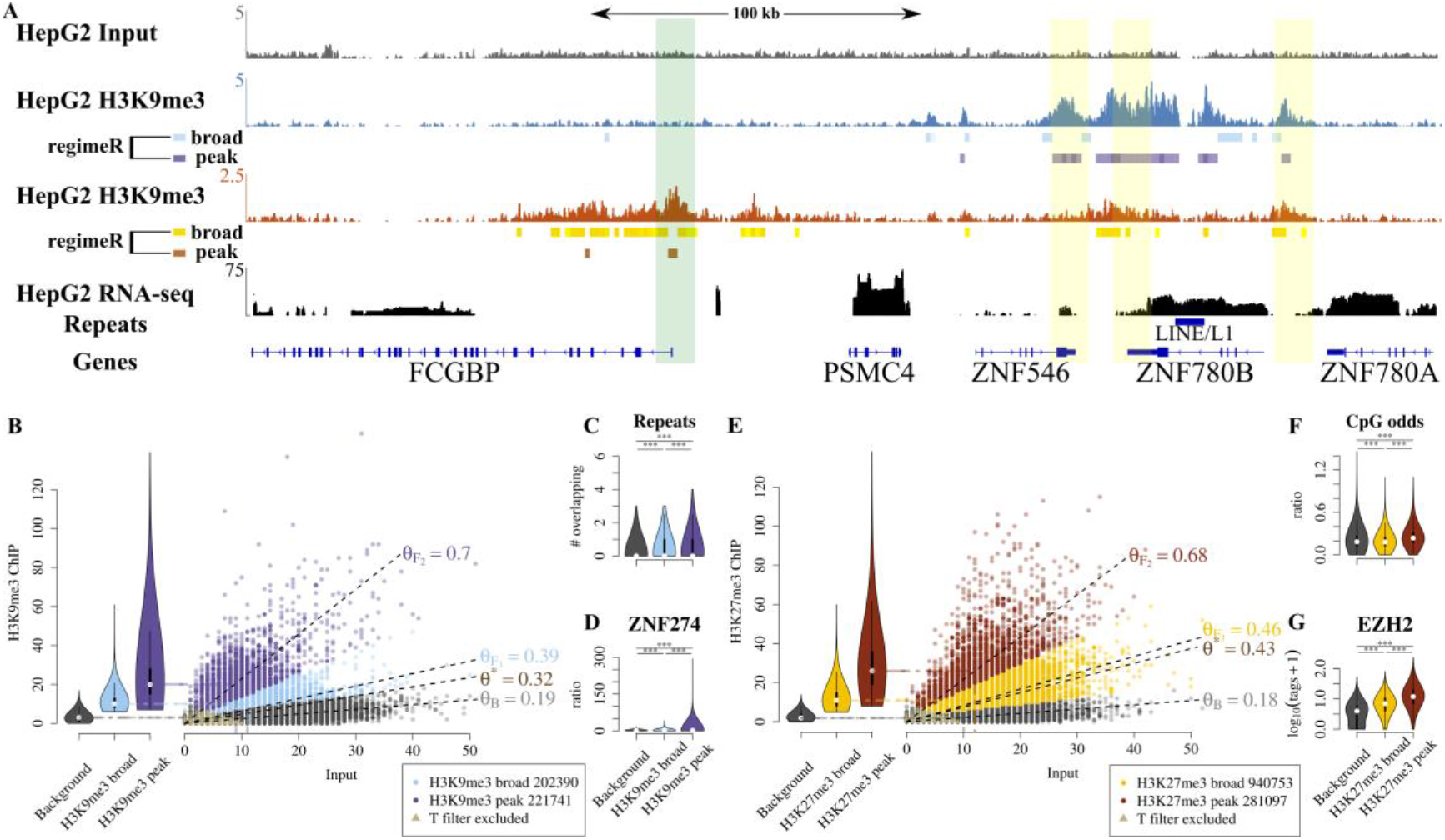
H3K27me3 and H3K9me3 Enrichment Regime Identification in HepG2 cells with regimeR. (A) Input (grey), H3K9me3 (blue), H3K27me3 (orange) and RNA-seq (black) coverage around a ZNF cluster on chromosome 19 in HepG2 cells. Individual regimeR-computed regimes are displayed as boxes below respective tracks. The 5’-ends of ZNF genes are marked with high H3K9me3 enrichment (yellow overlay) and the promoter of FCGBP is marked by a H3K27me3 peak within a broad H3K27me3 domain (green overlay). (B) regimeR identifies broad and peak H3K9me3 enrichment. (C-D) H3K9me3 peaks are significantly enriched for repeats (C) and ZNF274 ChIP-seq reads (D) as compared to both background and broad regions. (E) regimeR identifies broad and peak H3K27me3 enrichment. (F-G) H3K27me3 peaks have significantly greater CpG odds (F) and EZH2 binding (G) as compared to background and broad regions.

Some ChIP-seq peak callers perform worse when the sequencing depth in the ChIP library is reduced (Teytelman et al. 2013). To show the robustness of enrichR, we used the unified gold standard to benchmark all assessed tools on an *in silico* down sampled sequencing library (Supplemental Note). enrichR and MACS2 called >90% of the gold standard at 50% (30%) of the original H3K4me3 (H3K36me3) sequencing depth (Supplemental Fig. 4) suggesting that both methods are specific in even shallow sequenced ChIP libraries.

ChIP-seq coverage normalization based on *bona-fide* background regions is also done by NCIS (Liang and Keles 2012). For H3K36me3 NCIS estimated a normalization factor that was ~1.5-fold smaller than *θ\** and enrichR’s *θ*_*B*_ was ~2-fold smaller than *θ\** (Supplemental Fig. 5, Supplemental Table 3). Thus, enrichR achieved a normalization almost equivalent to NCIS, despite using a different model.

### Enrichment Regime Identification in H3K27me3 and H3K9me3 in HepG2 cells with regimeR

Hither to discussed was the applicability of normR to a well-studied problem: the discrimination of enrichment against background. Here, we turn to a problem for which we had found to best of our knowledge no precedent in the literature: the discrimination of moderate enrichment from high enrichment. We can easily address this problem by increasing the number of foreground components in normR from one single component to multiple components (Methods). We refer to this approach as regimeR: In the case of two foreground components, regimeR disriminates a *peak regime* (high enrichment) and a *broad regime* (moderate enrichment) over the background. We applied regimeR to H3K9me3 and H3K27me3 ChIP-seq data from the hepatocarcinoma cell line HepG2 over the control.

Figure 3A depicts a representative region on Human chromosome 19 harbouring active and repressed genes. regimeR segmented the ChIP-seq enrichment into broad and peak regions. For example, three H3K9me3 peaks flanked by moderate enrichment were detected by regimeR at the 3’-ends of ZNF546 and ZNF780A/B. Similarly, a H3K27me3-peak within a H3K27me3-broad domain was identified by regimeR at the “Fc Fragment Of IgG Binding Protein” gene promoter.

For H3K9me3, 14.7% of the HepG2 epigenome got classified into 202,390 broad (47.8%; 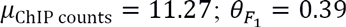) and 221,741 peak regions (52.2%; *μ*_ChIP counts_ = 23.75; 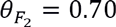; Figure 3B). Both H3K9me3-broad and –peak regions showed a statistically significant overlap with repetitive DNA elements (Wilcoxon-signed-rank test; P ≤ 0.001; Figure 3C, Supplemental Fig. 6A), which is a reported feature of H3K9me3 marked constitutive heterochromatin (Wang et al. 2016). Moreover, H3K9me3-peak regions showed significantly higher levels of ZNF274 than background and H3K9me3-broad regions (Wilcoxon-signed-rank test; P ≤ 0.001, Figure 3D), in line with the idea that ZNF274 recruits the H3K9 methyltransferase SETDB1 (Frietze et al. 2010). Thus H3K9me3-peak regions may coincide with nucleation sites for heterochromatin assembly at genomic repeat elements.

For H3K27me3, regimeR called 42.4% of the HepG2 epigenome H3K27me3-enriched (1,221,850 1kb regions) and subdivided this into 940,753 broad (77%, *μ*_ChIP counts_ = 12.03; 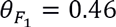) and 281,097 peak regions (23%, 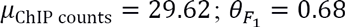 Figure 3E). H3K27me3 covered three times more of the genome than H3K9me3, yet, with a lower fraction of peak regions than in H3K9me3. Moreover, the vast majority H3K9me3 and H3K27me3 regimes were mutually exclusive in HepG2 cells (Supplemental Fig. 6B). H3K27me3-peak regions were characterized by a higher CpG odds ratio (CpG-content corrected for GC content) than both broad or background regions (Figure 3F, Supplemental Fig. 6C). In conjunction with an elevated conservation (Supplemental Fig. 6D) and a statistically significant overlap with annotated TSSs (Fisher’s signed exact test; P ≤ 0.001; odds ratio = 1.98; Supplemental Table 4) this reaffirms that the TSSs targeted for peak H3K27me3 levels are high CpG promoters (Saxonov et al. 2006). Similar to H3K9me3-peak regions, H3K27me3-peak regions were significantly enriched for the enzyme that catalyzes the modification, *i.e.* EZH2 (Müller et al. 2002; Kuzmichev et al. 2002; Cao et al. 2002; Czermin et al. 2002) (Wilcoxon signed-rank test; P ≤ 0.001, Figure 3G). Together these observations suggest that H3K27me3-broad and -peak regions show distinct characteristics with respect to CpG content, localization and EZH2 levels.

The observation that both H3K9me3- and H3K27me3-peak regions were associated with significantly higher levels of their catalyst than broad- and background regions indicates that they correspond to nucleation sites for heterochromatin assembly. In line with this observation we found that most H3K9me3-peak regions are either embedded in an H3K9me3 broad domain (43.4%) or at the border of a broad domain (35.1%). The vast majority of H3K27me3-peak regions were embedded in an H3K27me3 broad domain (82.8%) where both regimes showed elevated conservation (Supplemental Fig. 6D). On the contrary, H3K9me3-peaks were less conserved than broad regions further supporting aforementioned idea that repetitive elements recruit the H3K9me3 methyltransferase.

### Difference Calling in Primary Human Hepatocytes and HepG2 cells with diffR

In addition to discriminating enrichment from background, another important task consists of identifying epigenetic alterations between conditions, e.g. healthy versus diseased or between cell-types. normR can address this problem by calling differential enrichment between ChIP-seq experiments from two conditions, referred to as “diffR”. We applied diffR to H3K4me3 and H3K27me3 ChIP-seq data from primary human hepatocytes (PHH) and the hepatocarcinoma cell line HepG2 (Methods). We compared the diffR results to those obtained by calling mutually exclusive enrichment with enrichR on the two conditions separately, referred to as “enrichR-compare”. Additionally, we compared diffR results to three existing tools, namely ChIPDiff (Xu et al. 2008), histoneHMM (Heinig et al. 2015) and ODIN (Allhoff et al. 2014).

Visual inspection of a 50kb region on chromosome 19 confirmed that most H3K4me3/K27me3-enriched regions were common between HepG2 and PHH (Figure 4A). However, some enrichment was cell-type specific and was called by all methods, e.g. HepG2-specific H3K27me3-enrichment upstream of E2F2. However, differences in the histone modification level within mutually enriched regions were apparent, e.g. the increase in H3K4me3-enrichment at the E2F2 promoter in HepG2 could be identified by diffR, ChIPDiff and ODIN. E2F transcription factors are important regulators of the cell cycle (Sardet et al. 1995; Sylvestre et al. 2007; Ramboer et al. 2014). E2F2 is expressed in HepG2 but not in PHH suggesting that the induction of E2F2 might be linked to the much higher proliferative potential in HepG2 cells than in PHH. Further downstream of E2F2, enrichR identified a H3K27me3-differentially enriched domain accompanied by an emerging H3K4me3 peak in HepG2 cells. Thus, the induction of E2F2 in HepG2 may be explained by the opening of an enhancer at this region supported by reported binding of RNA polymerase 2 and CTCF in HepG2 cells (ENCODE Project Consortium 2012).

**Figure 4.**
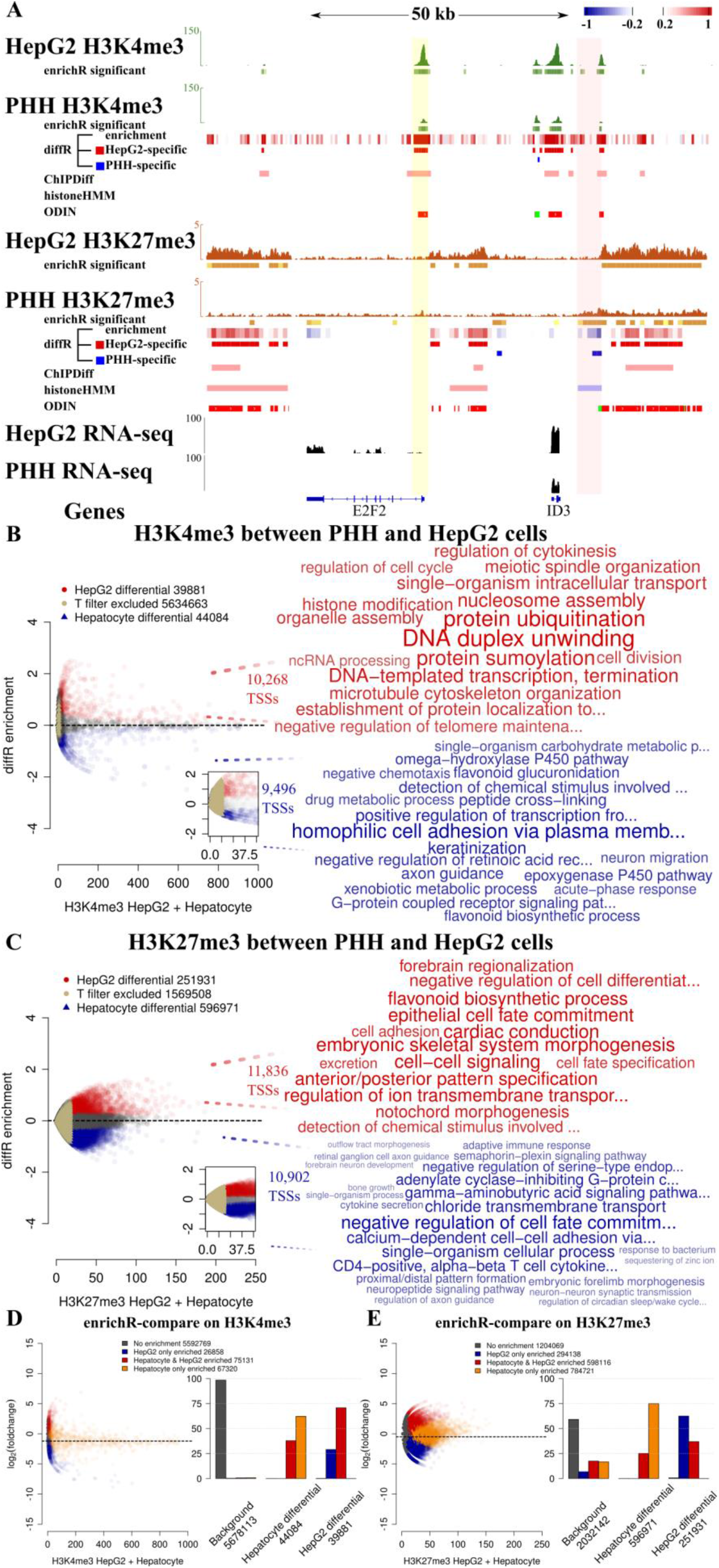
Difference Calling on H3K4me3 and H3K27me3 in Primary Human Hepatocytes (PHH) and HepG2 cells with diffR. (A) Input (grey), H3K4me3 (green), H3K27me3 (orange) and RNA-seq (black) coverage around E2F Transcription Factor 2 promoter (E2F2, yellow overlay) locus in Primary Human Hepatocytes (PHH) and HepG2 cells. A region ~40kb upstream of the E2F2 promoter shows significant differential enrichment for H3K4me3 and H3K27me3 (pink overlay). enrichR-computed enriched regions displayed as boxes below to respective. Differentially enriched regions are displayed as red (HepG2 conditional) or blue (PHH conditional) boxes for diffR, ChIPDiff, histoneHMM and ODIN. (B,C) diffR recovers conditional differences in H3K4me3 (B) and H3K27me3 (C) enrichments that cover transcriptional start sites (TSSs) driving genes functioning in cell metabolism and development (wordclouds right panel). (D,E) enrichR-compare identifies H3K4me3 (D) and H3K27me3 (E) mutually exclusive enrichment between PHH and HepG2 cells, but can not detect differences in histone modification level.(Right panels) diffR regions fall into enrichR-compare called regions of mutually exclusive enrichment but also resolve significant differences in ChIP-seq signal not detected by enrichR-compare.

For H3K4me3, diffR recovered 59,288 500bp regions (14Mb) as being differentially enriched between HepG2 and PHH (Figure 4B). Of these, 27,913 regions had a higher enrichment in HepG2 which overlapped 10,268 TSSs driving genes mainly related to the DNA replication and cell division. 31,375 PHH-specific H3K4me3 regions upregulated 9,496 TSSs of genes associated with liver function (P450 pathway) and tissue characteristics (keratinization, cell adhesion) absent in the HepG2 cell line. For H3K27me3, diffR reported 800,073 1kb regions (800Mb) as differentially H3K27me3-enriched (Figure 4C). Out of these 215,466 revealed HepG2-specific repression at 11,836 TSSs of genes regulating morphogenesis and cell-cell signaling. On the other hand, the 584,607 PHH-specific regions repressed 10,902 TSSs of genes functioning in cell fate commitment and immune response. Taken together, diffR uncovered functional differences related to immortalization of liver cells solely based on two ChIP-seq experiments.

Another normR approach can detect conditional differences by calling individual ChIP-seq enrichment over control for each condition and then identify mutually exclusive enrichment, referred to as “enrichR-compare”. We used this approach to benchmark results obtained from diffR. Genome-wide H3K4me3 enrichR-compare analysis revealed that most enriched 500bp regions were common in HepG2 and PHH (101,989, Figure 4D), while 26,858 were HepG2-and 67,320 PHH-specific. As expected, the comparison to enrichR-compare also revealed that by a majority diffR difference calls were either mutual exclusive enrichment or changes in the level of enrichment (Figure 4D, Supplemental Table 5). For H3K27me3, enrichR-compare revealed that most H3K27me3-enriched regions were common in HepG2 and PHH (892,254, Figure 4E), while 294,138 were HepG2- and 784,721 were PHH-specific. Again, diffR was very specific in capturing both mutual exclusive enrichment and changes in the level of enrichment (Figure 4E), However, we observed a discrepancy in sensitivity: 58.6% (44%) of the H3K4me3 (H3K27me3) mutually exclusive regions were not called by diffR leading to contradictory results (Supplemental Fig. 7A,B; Supplemental Table 5). Interestingly, most of the discrepancies were attributed to a more strict P-value filter to eliminate low power (i.e. low count) regions in the two-sided binomial test in diffR (Methods). By applying the diffR P-value filter to enrichR-compare, results became substantially more concordant, e.g. 2.99% (319) false negatives for H3K4me3 in HepG2 cells (Supplemental Fig. 7 C,D, Supplemental Table 5).

In addition, some discrepancies between diffR and enrichR-compare may be attributed to Copy Number Variations (CNVs) in HepG2 cells which are prevalent in immortalized cell types (Conrad et al. 2010; Shirley et al. 2012). To alleviate this problem we ran diffR on HepG2 and PHH Input tracks with 20 and 50kb windows (Supplemental Fig. 8). Assuming that there are no CNVs in the PHH data, diffR recovered 91% of 6,487 windows (odds-ratio=112.7) which overlap 80 annotated amplifications in HepG2 (ENCODE Project Consortium 2012) (13% of genome; median(*length*)=163kb). Nevertheless, diffR failed to detect 88% of 249 windows (odds-ratio=40.8) that overlap 170 annotated very short heterozygous and homozygous deletions (6% of genome; median(*length*)=9kb). Despite this, the discrepancies between enrichR-compare and diffR were partially removed when filtering results for diffR called CNVs (Supplemental Fig. 7E,F, Supplemental Table 5) to a similar extend than filtering for experimentally validated CNVs (Supplemental Fig.7G,H, Supplemental Table 5).

Next, we compared genome-wide diffR results to those obtained from ChIPDiff, histoneHMM and ODIN. To this end we once more defined a gold standard based among a consensus vote among the tools (Supplemental Note): ChIPDiff was most precise (*μ*_Precision_ = 0.70) and diffR had the highest recall (*μ*_Recall_ *=* 0.80) together with the best F1-scores (*μ*_F1-score_ = 0.50; Supplemental Table 6). A unified gold standard of all tool-specific gold standards revealed that most tool-specific regions were called by diffR (28.9Mb) and ODIN (25.4Mb) for H3K4me3 and by ODIN (701.7Mb) and histoneHMM (689.1Mb) for H3K27me3 (Supplemental Table 7). Turning to absolute fold changes, the unified gold standard showed highest levels together with diffR, ChIPDiff and histoneHMM (Supplemental Fig. 9A,B). In terms of read coverage, diffR- and ODIN-specific regions had highest counts (Supplemental Fig. 9C,D). In conclusion, diffR identified conditional differences for, both, H3K4me3 and H3K27me3 which were supported by a good classifier performance, a high absolute fold change as well as an inference-adequate read coverage eliminating low power regions.

## Discussion

In summary, we present an extendable methodology called “normR” that enables the extensive analysis of ChIP-seq data in epigenetic studies (Fig. 5). By modeling foreground and background jointly, normalization and enrichment calling are performed simultaneously. The implicit modeling of the effect of enrichment on the overall read statistics increases the sensitivity in detecting shallow differences in ChIP enrichment even in low signal-to-noise ratio data. Furthermore, we demonstrated the suitability of the normR approach for the identification of distinct epigenetic enrichment regimes in hepatocarcinoma cells and the quantification of conditional epigenetic differences between hepatocarcinoma cells and their tissue-of-origin. We envision how normR enrichment calling augments today’s epigenetic analyses ranging from clustering (Nair et al. 2014) to visualization Ramírez et al. 2014).

**Figure 5.**
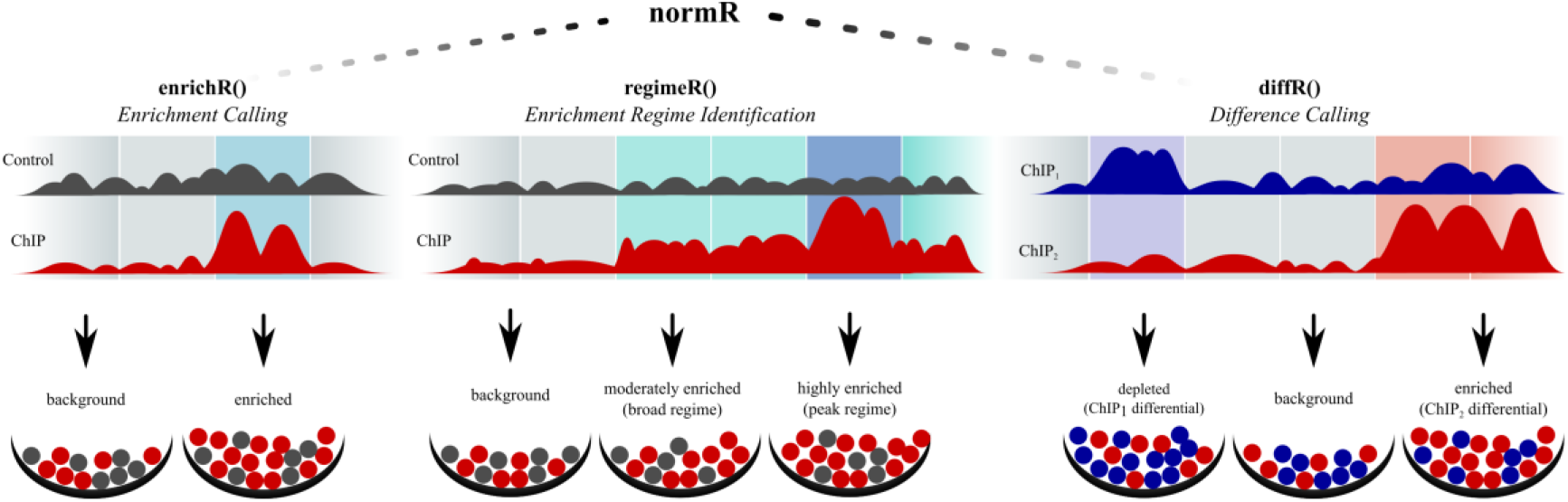
The normR Approach: A Robust and Broadly Applicable Methodology for Normalization and Difference Calling in ChIP-seq Data. The translational normR methodology allows for the calling of ChIP enrichment over a user-specified control, the identification of distinct ChIP enrichment regimes and the quantification of differences in ChIP signal level between two conditions.

Firstly we used normR to call enrichment in high (H3K4me3) and low (H3K36me3) signal-to-noise ratio ChIP-seq data, referred to as “enrichR”. Auxiliary information such as DNA methylation and expression supported the enrichR-based classification. Given the difficulty inherent in the ill-defined problem represented by ChIP-seq analysis we introduce a novel binary classifier statistic that defines a gold standard based on a consensus vote among seven published ChIP-seq peak callers. Our findings indicated that enrichR performs equally well as previously described approaches in ChIP-seq tracks with high signal-to-noise-ratio such as H3K4me3. Furthermore, enrichR outperformed existing tools in the detection of low levels of genuine enrichment in low signal-to-noise ratio data such as H3K36me3. We attribute the superior performance in the latter scenario to our sensitive normalization technique which accounts not only for varying sequencing depth but specifically addresses the effect of ChIP enrichment on the overall read statistics. The sensitive enrichR approach is an asset in future studies on epigenetic signatures and segmentations.

Secondly normR was used to facilitate the discrimination of peak- and broad-regions against background in a single analysis, referred to as “regimeR”. The analysis of H3K9me3 and H3K27me3 in HepG2 cells revealed that there exist distinct characteristics of peak- and broad regions in these heterochromatic marks. Specifically, H3K9me3 peaks were enriched for ZNF274 at repetitive elements. High enrichment of H3K9me3 at these sites can be explained by the recruitment of the H3K9 methyltransferase SETDB1 by ZNF274 (Frietze et al. 2010). H3K27me3 peaks were found within broad H3K27me3 domains at conserved CpG-dense regions bound by EZH2, supporting the idea of CpG-enriched polycomb recruitment sites (Tanay et al. 2007). Taken together, our regimeR-based study suggests that H3K9me3 and H3K27me3 peaks correspond to nucleation sites for heterochromatin assembly. In the future, regimeR will prove useful in studies of heterogeneity in cellular epigenetic markings to identify regions of promiscuous protein binding.

Finally we presented normR for the direct comparison of two ChIP-seq experiments, referred to as “diffR”. Our diffR-based comparison of H3K4me3 and heterochromatic H3K27me3 between HepG2 cells and PHH revealed conditional differences associated to cell function and immortalization, e.g. a potential E2F2 enhancer region made accessible in HepG2 cells. Interestingly, H3K27me3 covered a smaller fraction of the HepG2 genome as compared to PHH. Using a statistic of mutually exclusive enrichment by enrichR-compare and consensus votes among previously developed difference callers, we showed that diffR performs outstandingly in the detection of conditional differences in ChIP-seq data. Furthermore, we could show that diffR’s accuracy can be increased by incorporation of CNV information, as measured experimentally or by using diffR on two Input experiments. In the future, a more principled approach of the joint modelling of conditional ChIP-seq tracks together with their control is desirable.

Taken together normR proved as a versatile and sensitive toolbox for the discrimination of enrichment against background (“enrichR”), the unprecedented detection of enrichment regimes such as peaks and broad enrichment (“regimeR”) and the direct quantification of differences between two conditions (“diffR”). We anticipate that normR will be applied to all enrichment based sequencing technologies like MeDIP-seq and HiC. In fact, a derivate of normR has recently been used to identify co-localizing histone modifications in a novel reChIP-seq data set (Kinkley et al. 2016) where the background estimation is complicated by the presence of enrichment in the control experiment. In the future, an automated determination of the number of enrichment components in the normR model will be adjuvant in studying epigenomic heterogeneity in conjunction with recently reported single cell ChIP-seq data (Rotem et al. 2015).

## Methods

### The normR Methods

Given two vectors of integers *r* (control) and *s* (treatment) of identical length *n,* we model the read counts from the ChIP and control by a binomial *m*-mixture model:

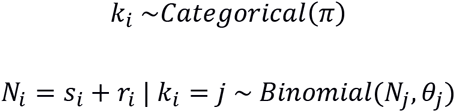

with *i* = 1,…,*n* and 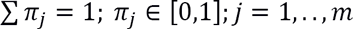. Given this model, normR follows a two step procedure: (i) The mixture model is fit by expectation maximization (EM; (Dempster et al. 1977)) using the likelihood function,

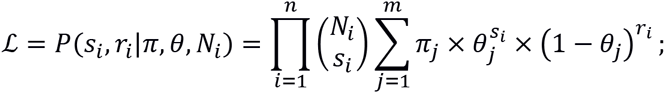

and (ii) each entry (*r*_*i*_, *s*_*i*_) is tested for significance against a fitted background to component to label enriched regions.

In a preprocessing stage, the vectors *r* and *s* are filtered for entries where *r = s* = 0 because no assertion about their enrichment state can be made. Secondly, a map of unique (*r*, *s*) tuples is created to reduce the number of computations needed which improves runtime substantially. In the first mode fitting step, the EM is initialized with *π* sampled from *U* (0,1) and *θ* sampled from *U*(0.001, *θ*\*). Upon convergence with *ɛ* ≤ 0.001, an enrichment factor (average fold enrichment) 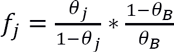 is computed for each mixture component j *≠* B, where *θ*_*B*_ the smallest of {*θ*_1_,…, *θ*_*m*_} (the closest to *θ\**) in the case of enrichment (difference) calling. The EM is run 10 times per default to find the fit with greatest 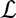. In the second step, every (*r*_*i*_, *s*_*i*_) is tested for significance against the background component. Resulting P-values are filtered using the T method (Dialsingh et al. 2015) (P-value threshold 0.0001 per default) to take into account the discreteness of P-values for a correct estimation of the proportion of true null hypotheses. T-filtered P-values are transformed to q-values for FDR correction (Storey 2002). Additionally, a normalized enrichment *e*_i_ is calculated for every entry (*r*, *s*)_*i*_ with 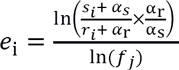 where 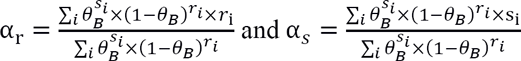 represent a model specific pseudo count for control and treatment, respectively. The normalized enrichment can be written to bigWig or bedGraph format for convenient display in a genome browser of choice, *e.g.* UCSC genome browser (Speir et al. 2016) or Integrative Genomics Viewer Robinson et al. 2011.

In the case of enrichment calling two components (background, enrichment) are fit with the enrichR subroutine of the normR package. Herein, the background model *θ*_*B*_ is set to the mixture component with smallest *θ*. For difference calling, three components (background, control enriched, treatment enriched) are fit with the diffR subroutine for *r* (condition 1) and s (condition 2) counts. The background model is set to *θ*_*B*_ closest to θ*. The diffR T method uses the maximal threshold estimated from P-values for θ_*B*_ fit for either (*r, s*) or the label-switched (*s*, *r*). For regime calling, the regimeR subroutine fits an arbitrary number of components representing background plus a fixed number of enrichment regimes. Identically to enrichment calling, the background model is set to the mixture component with smallest *θ*. In a second step, every significantly enriched bin passing the P-value filter (see above) is assigned to an enrichment regime by *Maximum A Posteriori*.

Note that by nature the binomial mixture model assumes the independence between regions which is valid for a sufficiently large bin size (*i.e*. fragment length). Consequently, the usage of a binomial mixture model improves computational runtime. The normR algorithm is implemented in C++ and R. A ready-to-use R-package can be obtained from http://bioconductor.org/packages/normr where also a tutorial on use cases can be found.

### ChIP-seq in primary human hepatocytes and HepG2 cells

HepG2 cells and human hepatocytes, obtained from donors after written consent by tissue resection and perfusion (Godoy et al. 2013), have been fixed in for 5 minutes in 1% formaldehyde. Formaldehyde has been quenched using 125 mM glycine and cells have been washed in PBS, pelleted and snap-frozen in liquid nitrogen. Five (human hepatocytes) to ten (HepG2) million cells have been processed for chromatin preparation, using the NEXSON protocol, as previously described (Arrigoni et al. 2016). After chromatin sonication, samples have been quality controlled to check chromatin recovery and fragment size distribution as previously described.

Prior ChIP, chromatin has been diluted 1:2 in the ChIP buffer H from the Diagenode Auto histone ChIP-seq kit (C01010022), supplemented with protease inhibitor cocktail. Chromatin from 100,000 to 500,000 cells has been incubated with one microgram of the following antibodies: H3K4me3 (C15410003), H3K36me3 (C15410192), H3K9me3 (C15410193), H3K27me3 (C15410195), all from Diagenode. ChIP has been performed using the automated platform SX-8G IP-Star (Diagenode), with the following parameters: “indirect ChIP”, 200 µl ChIP volume, 14 hours of antibody incubation, 4 hours of beads incubation, and 5 minutes beads washes. After the DNA elution from the beads, samples were collected, RNaseA-treated, de-proteinized and decrosslinked overnight at 65 °C. Input samples have been prepared by taking 1% of the starting chromatin before ChIP and by decrosslinking it together with the ChIP samples. DNA has been manually purified using the Qiagen minElute columns.

Libraries from 2 to 10 ng of purified DNA have been prepared using the NEBNext Ultra DNA library preparation kit (NEB, E7370S) following manufacturer’s instruction and skipping the size selection. Libraries have been sequenced paired-end, with a read length of 50 bp, on an Illumina HiSeq 2500 (version 3 chemistry).

### RNA-seq in primary human hepatocytes and HepG2 cells

Trizol extration was used for preparation of Total RNA according to the manufacturer’s guidelines and as described in (Lappalainen et al. 2013). An Agilent Bioanalyzer (Agilent, Santa Clara, USA) was used to check RNA integrity following the manufacturer’s guidelines.

Strand-specific sequencing libraries for mRNA and total-RNA were constructed for the HepG2 cells and human hepatocytes using the TruSeq stranded Total RNA kit (Illumina Inc, San Diego, USA) starting from 500 ng of the total RNA of the samples. Illumina HiSeq2000 was used to perform the sequencing (101-nucleotide paired-end reads for each library) resulting in the creation of about 100 million reads per library.

The reads were aligned to the NCBI 37.1 version of human genome using TopHat v2.0.11 (Kim et al. 2013) in the settings “--library-type fr-firststrand” and “--b2-very-sensitive”.

Reads mapping to genes were counted using htseq-count from HTSeq-0.6.1p1 (Anders et al. 2015) in '-f bam -s reverse -m union -a 20' setting. Annotation file for running htseq-count was downloaded from GENCODE release 19 (GRCh37.p13).

### Quantification of reads

Paired-end reads from Input, H3K4me3, H3K27me3, H3K36me3 and H3K9me3 ChIP-seq for primary human hepatocytes and HepG2 cells were mapped with bwa (version 0.6.2) against hg19. Fragment coverage tracks for browser display were generated with deepTools (Ramírez et al. 2016) in 25 bp windows (-bs 25) considering only first reads in a properly mapped pair (--samFlag 66) with a mapping quality of at least 20 (--MinMappingQuality 20) and normalized to the effective genome size (--normalizeTo1x 2451960000):

~~~
bamCoverage -bam in.bam -o out.bw -of bigwig -bs 25 \
  --samFlag 66 --minMappingQuality 20 --normalizeTo1x 2451960000
~~~

For enrichment and peak calling, only regions on regular autosomes (chr1-chr22; 2.9Gb) were used:

~~~
require(GenomeInfoDb)
genome <- fetchExtendedChromInfoFromUCSC(“hg19”)
genome <- genome[which(!genome$circular &
   genome$SequenceRole=="assembled-molecule"), 1:2]
genome <- genome[grep("X|Y|M", genome[, 1], invert=T),]
require(GenomicRanges)
genome.gr <- GRanges(
 seqnames = genome[, 1],
 ranges = IRanges(start = 1, end = genome[, 2]),
 seqinfo = Seqinfo(
  seqnames = genome[,1],
  seqlengths = genome[,2],
  genome = “hg19”)))
}
~~~

For paired end data, we considered only reads with a mapping quality of at least 20 (mapqual=20). We regarded midpoints of properly mapped fragments (midpoint = TRUE) that were non-duplicated (filteredFlag=1024) and within 100 to 220 bp in length (tlenFilter=c(100,220)) in 500 (1,000) bp windows for H3K4me3 (H3K27me3/K36me3/K9me3) with normR’s countConfigPairedEnd function:

~~~
require(normr)
countConfig <- countConfigPairedEnd(
 binsize = 500, #1000
 mapqual = 20,
 midpoint = TRUE,
 filteredFlag = 1024,
 tlenFilter = c(100,220)
 shift = 0
)
~~~

HepG2 CAGE data was downloaded from GSM849335 (Djebali et al. 2012). Primary human hepatocyte CAGE data was downloaded from CAGE http://fantom.gsc.riken.jp/5/datafiles/latest/basic/human.primary_cell.hCAGE/Hepatocyte%252c%2520donor2.CNhs12349.11603-120I1.hg19.nobarcode.bam (Fantom5 (FANTOM Consortium and the RIKEN PMI and CLST (DGT) et al. 2014)) Reads with mapping quality of at least 20 were counted with bamsignals (http://bioconductor.org/packages/bamsignals):

~~~
require(bamsignals)
cage <- bamProfile(
 bampath = “Cage.bam”,
 gr = genome.gr,
 binsize = 500, #1000
 mapqual = 20
)
~~~

EZH2 ChIP-seq alignments (GSM1003576) and the respective control alignment (GSM733780) were downloaded from the UCSC encode repository ((ENCODE Project Consortium 2012) hgdownload.cse.ucsc.edu/goldenPath/hg19/encodeDCC/wgEncodeBroadHistone/). For these single end data, we shifted reads by 100 bp in 3’ direction (shift=100) and counted in 500 (1,000) bp bins:

~~~
countConfig <- countConfigSingleEnd(
 binsize = 500, #1000
 mapqual = 20,
 filteredFlag = 1024,
 shift = 100
)
~~~

### Enrichment calling with enrichR

Read counts in H3K4me3 and H3K36me3 were modeled with 2 components in enrichR and the fitted background components were used for significance tests. Bins with q-value ≤ 0.05 (H3K4me3) and q-value ≤ 0.1 (H3K27me3/K36me3/K9me3) were called enriched and exported to bed tracks for display:

~~~
enrichment <- enrichR(
 treatment = “ChIP.bam”,
 control = “Input.bam”,
 genome = genome,
 countConfig = countConfig,
 procs = 24
)
exportR(
 x = enrichment,
 filename = “enriched.bed”,
 type = “bed”,
 fdr = 0.05 #0.1
)
~~~

### DNA-methylation in primary human hepatocytes and HepG2 cells

For whole-genome bisulfite sequencing we produced two types of NGS libraries to achieve even read coverage. Firstly, we used 100ng of DNA with the TruSeq DNA methylation kit (Illumina, San Diego, USA) according to the manufacturer’s protocol. The second type was done as previously described (Kinkley et al. 2016). Briefly, 2 µg of DNA were sheared using a Bioruptor NGS device (Diagenode, Liege, Belgium) and cleaned-up using Ampure beads XP (Beckman Coulter, Brea, USA). Then samples were subjected to end-repair, A-tailing and adaptor ligation steps using components of the TruSeq DNA PCR-Free Library Preparation Kit (Illumina). After bisulfite conversion involving the Zymo Gold kit (Zymo, Irvine, USA) the libraries were PCR amplified for 10-12 cycles. The amplified libraries were purified using Ampure beads XP and sequenced on three lanes of V3 paired-end flow cells (2x 100bp). Reads were mapped using BWA (Li and Durbin 2010)and methylation levels were called with Bis-SNP37 (Liu et al. 2011).

Beta values were calculated for each bin and weighted by coverage and number of CpGs *M* in that region: 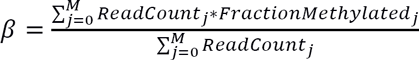. Only regions with at least 2 CpGs covered by reads were reported.

### Transcription Start Site Definition

54,763 promoters (extend 750bp down- and upstream of TSS) of 54,849 GENCODE genes (Harrow et al. 2012) obtained by using GenomicFeatures R package (Lawrence et al. 2013):

~~~
require(GenomicFeatures)
gencode <- loadDb("data/gencode.v19.annotation.transcriptDb.sqlite")
genes <- genes(gencode)
proms <- unique(promoters(genes, upstream=750, downstream=750))
~~~

### MACS, DFilter, CisGenome, SPP, BCP and MUSIC Peak Calling

Peaks were called with MACS2 (Feng et al. 2012) (v2.1.0.20150731), DFilter (Kumar et al. 2013) (v1.6), CisGenome (Ji et al. 2008), SPP (Kharchenko et al. 2008), BCP Xing et al. 2012) (v1.1) and MUSIC (Harmanci et al. 2014). A FDR threshold of 0.1 was used. To compare called peaks by above methods to enrichR called regions, overlap of peaks with 500 bp (1,000 bp) windows was calculated for H3K4me3 (H3K36me3). See Supplemental Note for details.

### Normalization Factor Comparison with NCIS

NCIS (Liang and Keles 2012) was run in R to calculate the normalization factor for comparison to enrichR’s normalization factor:

~~~
require(NCIS)
ncis <- NCIS(
 chip.data = “ChIP.bed”,
 input.data = “Control.bed”,
 data.type = "BED",
 chr.vec = genome[,1],
 chr.len.vec = genome[,2]
)
ncis.norm <- ncis$est
~~~

### Regime calling with regimeR

Read counts in H3K27me3 and H3K9me3 in HepG2 cells were modeled in regimeR with 3 components (background, moderate enrichment, high enrichment). Bins with FDR q-value ≤ 0.1 were called enriched and assigned to an enrichment component by *Maximum A Posteriori* and exported to bed using normR’s exportR function:

~~~
regimes <- regimeR(
 treatment = “ChIP.bam”,
 control = “Input.bam”,
 genome = genome,
 models = 3,
 countConfig = countConfig,
 procs = 24
)
exportR(
 x = regimes,
 filename = “regimes.bed”,
 type = “bed”,
 fdr = 0.05 #0.1
)
~~~

### Difference calling with diffR

Read counts in H3K4me3 and H3K27me3 in primary human hepatocytes (control) and HepG2 cells (treatment) were modeled in diffR with 3 components (background/no difference, treatment-enriched, control-enriched) and the mixture component with θ_*j*_ closest to θ* was used as background for a two-sided significance test. Bins with q-value ≤ 0.05 (0.1) for H3K4me3 (H3K27me3) were called differentially enriched and assigned to treatment or control by *Maximum A Posteriori*.

~~~
diffs <- diffR(
 treatment = “ChIP1.bam”,
 control = “ChIP2.bam”,
 genome = genome,
 countConfig = countConfig,
 procs = 24
)
exportR(
 x = diffs,
 filename = “differences.bed”,
 type = “bed”,
 fdr = 0.05 #0.1
)
~~~

To analyze differentially enriched regions for precision and recall, *mutually exclusive enrichment* in control (treatment) was obtained by considering enrichR() calls present only in control (treatment) with respect to treatment (control). For a fair comparison, only significant regions with a posterior of ≥ 0.50 were considered.

### Gene Ontology Analysis

We used topGO (Aibar et al. 2015) on gene ontology “Biological Process” (BP) with algorithms “classic” (algorithm=”classic”) and “elim” (algorithm=”elim”) for statistics “fisher” (statistic=”fisher”) and “ks” (statistic=”ks”) for GENCODE gene IDs mapped to Ensembl gene IDs. The “ks” statistic allows for supplying a score for each entity. We used the diffR calculated q-value as score. We retained only top 1,000 (n=1000) GO terms ordered by “elim” algorithm and ranked by “classic” algorithm calculated P-values:

~~~
require(topGO)
#get GO annotated Ensembl Genes
go2ensembl <- annFUN.org(ontology, mapping="org.Hs.eg.db", ID="ensembl")
#get GENCODE genes and filter these for the ones in gene universe
gencode <- loadDb("data/gencode.v19.annotation.transcriptDb.sqlite")
gene.universe <- intersect(
 unique(GenomicFeatures::genes(gencode)$genes),
 unique(unlist(go2ensembl))
)
#set diffR pvalue as score for differentially modified TSSs
idx <- gene.universe %in% diffTSSs
allGenes <- 1-as.integer(idx)
names(allGenes) <- gene.universe
allGenes[idx] <- pvals[diffTSSs %in% gene.universe]
goData <- new("topGOdata",
 description="diffR differential TSS histone marking study (scored)",
 ontology=”BP”,
 allGenes=allGenes, geneSel=function(p) { return(p <= 0.05) },
 annot=annFUN.GO2genes, GO2genes=go2ensembl, #GO mapping for ensembl IDs
 nodeSize=10
)
#testing
resultFisher <- runTest(goData, algorithm="classic", statistic="fisher")
resultKS <- runTest(goData, algorithm="classic", statistic="ks")
resultKS.elim <- runTest(goData, algorithm="elim", statistic="ks")
#compile results
resDf <- GenTable(goData,
 classicFisher = resultFisher,
 classicKS = resultKS,
 elimKS = resultKS.elim,
 orderBy ="elimKS",
 ranksOf = "classicFisher",
 topNodes=1000
)
~~~

### ChIPDiff, histoneHMM and ODIN Difference

Differences for H3K4me3 (H3K27me3) between Hepatocytes and HepG2 cells were called with ChIPDiff (Xu et al. 2008), histoneHMM (v1.6) (Heinig et al. 2015) and ODIN (v0.4) (Allhoff et al. 2014). A FDR threshold of 0.1 was used. To compare called peaks by above methods to diffR called regions, overlap of peaks with 500 bp (1,000 bp) windows was calculated for H3K4me3 (H3K27me3). See Supplemental Note for details.

### HepG2 Genotyping

HepG2 genotype information for hg19 was generated by ENCODE/HudsonAlpha (GSM999286) and downloaded from UCSC (http://hgdownload.cse.ucsc.edu/goldenPath/hg19/encodeDCC/wgEncodeHaibGenotype/wgEncodeHaibGenotypeHepg2RegionsRep1.bedLogR.gz).

## Data Access

H3K4me3, H3K9me3, H3K27me3, H3K36me3 ChIP-seq and Input data for primary human hepatocytes have been deposited at the “European Genome-Phenome Archive” under the accession EGAS00001002080. H3K4me3, H3K9me3, H3K27me3, H3K36me3 ChIP-seq and Input data for HepG2 have been deposited at the European Nucleotide Archive under the accession PRJEB7356.

## Acknowledgements

We thank the DEEP consortium for their extended help and support. This work was supported by the Bundesministerium für Bildung und Forschung ‘Deutsches Epigenom Programm’ [01KU1216C] (PR, JGH, JW, TM and HRC).

## Author Contributions

JH and HRC developed the methodology. JH implemented the algorithm, deployed the package and performed analyses. KG and CC provided the primary human hepatocyte samples. LA and NL performed ChIP-seq experiments. GG performed whole genome bisulfite sequencing. AS performed RNA-seq experiments. JGH, PR, JW, TM and HRC supervised the experiments and analyses and acquired funding. JH and HRC wrote with the help of TM the manuscript. All authors read and approved the manuscript.

## Disclosure Declaration

### Ethics approval and consent to participate

T.B.D.

### Consent for publication

T.B.D.

## References

Aibar S, Fontanillo C, Droste C, Las Rivas De J. 2015. Functional Gene Networks: R/Bioc package to generate and analyse gene networks derived from functional enrichment and clustering. Bioinformatics 31: 1686–1688.

Allhoff M, Seré K, Chauvistré H, Lin Q, Zenke M, Costa IG. 2014. Detecting differential peaks in ChIP-seq signals with ODIN. Bioinformatics 30: 3467–3475.

Anders S, Pyl PT, Huber W. 2015. HTSeq--a Python framework to work with high-throughput sequencing data. Bioinformatics 31: 166–169.

Arrigoni L, Richter AS, Betancourt E, Bruder K, Diehl S, Manke T, Bönisch U. 2016. Standardizing chromatin research: a simple and universal method for ChIP-seq. Nucleic Acids Res 44: e67.

Baubec T, Colombo DF, Wirbelauer C, Schmidt J, Burger L, Krebs AR, Akalin A, Schübeler D. 2015. Genomic profiling of DNA methyltransferases reveals a role for DNMT3B in genic methylation. Nature 520: 243–247.

Bonn S, Zinzen RP, Girardot C, Gustafson EH, Perez-Gonzalez A, Delhomme N, Ghavi-Helm Y, Wilczyński B, Riddell A, Furlong EEM. 2012. Tissue-specific analysis of chromatin state identifies temporal signatures of enhancer activity during embryonic development. Nat Genet 44: 148–156.

Cao R, Wang L, Wang H, Xia L, Erdjument-Bromage H, Tempst P, Jones RS, Zhang Y. 2002. Role of histone H3 lysine 27 methylation in Polycomb-group silencing. Science 298: 1039–1043.

Conrad DF, Pinto D, Redon R, Feuk L, Gokcumen O, Zhang Y, Aerts J, Andrews TD, Barnes C, Campbell P, et al. 2010. Origins and functional impact of copy number variation in the human genome. Nature 464: 704–712.

Czermin B, Melfi R, McCabe D, Seitz V, Imhof A, Pirrotta V. 2002. Drosophila enhancer of Zeste/ESC complexes have a histone H3 methyltransferase activity that marks chromosomal Polycomb sites. Cell 111: 185–196.

Dempster AP, Laird NM, Rubin DB. 1977. Maximum Likelihood from Incomplete Data via the EM Algorithm on JSTOR. Journal of the royal statistical society ….

Dialsingh I, Austin SR, Altman NS. 2015. Estimating the proportion of true null hypotheses when the statistics are discrete. Bioinformatics 31: 2303–2309.

Diaz A, Park K, Lim DA, Song JS. 2012. Normalization, bias correction, and peak calling for ChIP-seq. Stat Appl Genet Mol Biol 11: Article 9.

Djebali S, Davis CA, Merkel A, Dobin A, Lassmann T, Mortazavi A, Tanzer A, Lagarde J, Lin W, Schlesinger F, et al. 2012. Landscape of transcription in human cells. Nature 489: 101–108.

Eisenberg E, Levanon EY. 2013. Human housekeeping genes, revisited. Trends Genet 29: 569–574.

ENCODE Project Consortium. 2012. An integrated encyclopedia of DNA elements in the human genome. Nature 489: 57–74.

Ernst J, Kellis M. 2012. ChromHMM: automating chromatin-state discovery and characterization. Nat Methods 9: 215–216.

FANTOM Consortium and the RIKEN PMI and CLST (DGT), Forrest ARR, Kawaji H, Rehli M, Baillie JK, de Hoon MJL, Haberle V, Lassmann T, Kulakovskiy IV, Lizio M, et al. 2014. A promoter-level mammalian expression atlas. Nature 507: 462–470.

Feng J, Liu T, Qin B, Zhang Y, Liu XS. 2012. Identifying ChIP-seq enrichment using MACS. Nat Protoc 7: 1728–1740.

Flensburg C, Kinkel SA, Keniry A, Blewitt ME, Oshlack A. 2014. A comparison of control samples for ChIP-seq of histone modifications. Front Genet 5: 329.

Frietze S, O'Geen H, Blahnik KR, Jin VX, Farnham PJ. 2010. ZNF274 recruits the histone methyltransferase SETDB1 to the 3' ends of ZNF genes. PLoS ONE 5: e15082.

Godoy P, Hewitt NJ, Albrecht U, Andersen ME, Ansari N, Bhattacharya S, Bode JG, Bolleyn J, Borner C, Böttger J, et al. 2013. Recent advances in 2D and 3D in vitro systems using primary hepatocytes, alternative hepatocyte sources and non-parenchymal liver cells and their use in investigating mechanisms of hepatotoxicity, cell signaling and ADME. Arch Toxicol 87: 1315–1530.

Harmanci A, Rozowsky J, Gerstein M. 2014. MUSIC: identification of enriched regions in ChIP-Seq experiments using a mappability-corrected multiscale signal processing framework. Genome Biol 15: 474.

Harrow J, Frankish A, Gonzalez JM, Tapanari E, Diekhans M, Kokocinski F, Aken BL, Barrell D, Zadissa A, Searle S, et al. 2012. GENCODE: the reference human genome annotation for The ENCODE Project. Genome Res 22: 1760–1774.

Heinig M, Colomé-Tatché M, Taudt A, Rintisch C, Schafer S, Pravenec M, Hubner N, Vingron M, Johannes F. 2015. histoneHMM: Differential analysis of histone modifications with broad genomic footprints. BMC Bioinformatics 16: 60.

Heintzman ND, Hon GC, Hawkins RD, Kheradpour P, Stark A, Harp LF, Ye Z, Lee LK, Stuart RK, Ching CW, et al. 2009. Histone modifications at human enhancers reflect global cell-type-specific gene expression. Nature 459: 108–112.

Heintzman ND, Stuart RK, Hon G, Fu Y, Ching CW, Hawkins RD, Barrera LO, Van Calcar S, Qu C, Ching KA, et al. 2007. Distinct and predictive chromatin signatures of transcriptional promoters and enhancers in the human genome. Nat Genet 39: 311–318.

Hu J-L, Zhou BO, Zhang R-R, Zhang K-L, Zhou J-Q, Xu G-L. 2009. The N-terminus of histone H3 is required for de novo DNA methylation in chromatin. Proc Natl Acad Sci USA 106: 22187–22192.

Jain D, Baldi S, Zabel A, Straub T, Becker PB. 2015. Active promoters give rise to false positive “Phantom Peaks” in ChIP-seq experiments. Nucleic Acids Res 43: 6959–6968.

Ji H, Jiang H, Ma W, Johnson DS, Myers RM, Wong WH. 2008. An integrated software system for analyzing ChIP-chip and ChIP-seq data. Nat Biotech 26: 1293–1300.

Johnson DS, Mortazavi A, Myers RM, Wold B. 2007. Genome-wide mapping of in vivo protein-DNA interactions. Science 316: 1497–1502.

Karlić R, Chung H-R, Lasserre J, Vlahoviček K, Vingron M. 2010. Histone modification levels are predictive for gene expression. Proc Natl Acad Sci USA 107: 2926–2931.

Kharchenko PV, Tolstorukov MY, Park PJ. 2008. Design and analysis of ChIP-seq experiments for DNA-binding proteins. Nat Biotechnol 26: 1351–1359.

Kim A, Kiefer CM, Dean A. 2007. Distinctive signatures of histone methylation in transcribed coding and noncoding human beta-globin sequences. Mol Cell Biol 27: 1271–1279.

Kim D, Pertea G, Trapnell C, Pimentel H, Kelley R. 2013. TopHat2: accurate alignment of transcriptomes in the presence of insertions, deletions and gene fusions. Genome ….

Kinkley S, Helmuth J, Polansky JK, Dunkel I, Gasparoni G, Fröhler S, Chen W, Walter J, Hamann A, Chung H-R. 2016. reChIP-seq reveals widespread bivalency of H3K4me3 and H3K27me3 in CD4(+) memory T cells. Nat Commun 7: 12514.

Kumar V, Muratani M, Rayan NA, Kraus P, Lufkin T, Ng HH, Prabhakar S. 2013. Uniform, optimal signal processing of mapped deep-sequencing data. Nat Biotechnol 31: 615–622.

Kuzmichev A, Nishioka K, Erdjument-Bromage H, Tempst P, Reinberg D. 2002. Histone methyltransferase activity associated with a human multiprotein complex containing the Enhancer of Zeste protein. Genes Dev 16: 2893–2905.

Lappalainen T, Sammeth M, Friedländer MR, 't Hoen PAC, Monlong J, Rivas MA, Gonzàlez-Porta M, Kurbatova N, Griebel T, Ferreira PG, et al. 2013. Transcriptome and genome sequencing uncovers functional variation in humans. Nature 501: 506–511.

Lawrence M, Huber W, Pagès H, Aboyoun P, Carlson M, Gentleman R, Morgan MT, Carey VJ. 2013. Software for computing and annotating genomic ranges. PLoS Comput Biol 9: e1003118.

Li H, Durbin R. 2010. Fast and accurate long-read alignment with Burrows-Wheeler transform. Bioinformatics 26: 589–595.

Liang K, Keles S. 2012. Normalization of ChIP-seq data with control. BMC Bioinformatics 13: 199.

Liu Y, Siegmund KD, Laird PW, Berman BP. 2011. Bis-SNP: combined DNA methylation and SNP calling for Bisulfite-seq data. Genome Biol 13: R61–R61.

Long HK, Sims D, Heger A, Blackledge NP, Kutter C, Wright ML, Grützner F, Odom DT, Patient R, Ponting CP, et al. 2013. Epigenetic conservation at gene regulatory elements revealed by non-methylated DNA profiling in seven vertebrates. Elife 2: e00348.

Mammana A, Chung H-R. 2015. Chromatin segmentation based on a probabilistic model for read counts explains a large portion of the epigenome. Genome Biol 16: 151.

Meyer CA, Liu XS. 2014. Identifying and mitigating bias in next-generation sequencing methods for chromatin biology. Nat Rev Genet 15: 709–721.

Müller J, Hart CM, Francis NJ, Vargas ML, Sengupta A, Wild B, Miller EL, O'Connor MB, Kingston RE, Simon JA. 2002. Histone methyltransferase activity of a Drosophila Polycomb group repressor complex. Cell 111: 197–208.

Nair NU, Kumar S, Moret BME, Bucher P. 2014. Probabilistic partitioning methods to find significant patterns in ChIP-Seq data. Bioinformatics 30: 2406–2413.

Ooi SKT, Qiu C, Bernstein E, Li K, Jia D, Yang Z, Erdjument-Bromage H, Tempst P, Lin S-P, Allis CD, et al. 2007. DNMT3L connects unmethylated lysine 4 of histone H3 to de novo methylation of DNA. Nature 448: 714–717.

Perner J, Lasserre J, Kinkley S, Vingron M, Chung H-R. 2014. Inference of interactions between chromatin modifiers and histone modifications: from ChIP-Seq data to chromatin-signaling. Nucleic Acids Res 42: 13689–13695.

Ramboer E, De Craene B, De Kock J, Vanhaecke T, Berx G, Rogiers V, Vinken M. 2014. Strategies for immortalization of primary hepatocytes. J Hepatol 61: 925–943.

Ramírez F, Dündar F, Diehl S, Grüning BA, Manke T. 2014. deepTools: a flexible platform for exploring deep-sequencing data. Nucleic Acids Res 42: W187–91.

Ramírez F, Ryan DP, Grüning B, Bhardwaj V, Kilpert F, Richter AS, Heyne S, Dündar F, Manke T. 2016. deepTools2: a next generation web server for deep-sequencing data analysis. Nucleic Acids Res 44: W160–5.

Roadmap Epigenomics Consortium, Kundaje A, Meuleman W, Ernst J, Bilenky M, Yen A, Heravi-Moussavi A, Kheradpour P, Zhang Z, Wang J, et al. 2015. Integrative analysis of 111 reference human epigenomes. Nature 518: 317–330.

Robinson JT, Thorvaldsdóttir H, Winckler W, Guttman M, Lander ES, Getz G, Mesirov JP. 2011. Integrative genomics viewer. Nat Biotechnol 29: 24–26.

Rotem A, Ram O, Shoresh N, Sperling RA, Goren A, Weitz DA, Bernstein BE. 2015. Single-cell ChIP-seq reveals cell subpopulations defined by chromatin state. Nat Biotechnol.

Sardet C, Vidal M, Cobrinik D, Geng Y, Onufryk C, Chen A, Weinberg RA. 1995. E2F-4 and E2F-5, two members of the E2F family, are expressed in the early phases of the cell cycle. Proc Natl Acad Sci USA 92: 2403–2407.

Saxonov S, Berg P, Brutlag DL. 2006. A genome-wide analysis of CpG dinucleotides in the human genome distinguishes two distinct classes of promoters. Proc Natl Acad Sci USA 103: 1412–1417.

Shirley MD, Baugher JD, Stevens EL, Tang Z, Gerry N, Beiswanger CM, Berlin DS, Pevsner J. 2012. Chromosomal variation in lymphoblastoid cell lines. Hum Mutat 33: 1075–1086.

Sims D, Sudbery I, Ilott NE, Heger A, Ponting CP. 2014. Sequencing depth and coverage: key considerations in genomic analyses. Nat Rev Genet 15: 121–132.

Speir ML, Zweig AS, Rosenbloom KR, Raney BJ, Paten B, Nejad P, Lee BT, Learned K, Karolchik D, Hinrichs AS, et al. 2016. The UCSC Genome Browser database: 2016 update. Nucleic Acids Res 44: D717–25.

Storey JD. 2002. A direct approach to false discovery rates. Journal of the Royal Statistical Society: Series B (Statistical Methodology) 64: 479–498.

Sylvestre Y, De Guire V, Querido E, Mukhopadhyay UK, Bourdeau V, Major F, Ferbeyre G, Chartrand P. 2007. An E2F/miR-20a autoregulatory feedback loop. J Biol Chem 282: 2135–2143.

Tanay A, O'Donnell AH, Damelin M, Bestor TH. 2007. Hyperconserved CpG domains underlie Polycomb-binding sites. Proc Natl Acad Sci USA 104: 5521–5526.

Team RC. 2015. R: A Language and Environment for Statistical Computing. R Foundation for Statistical Computing, Vienna, Austria.

Teytelman L, Thurtle DM, Rine J, van Oudenaarden A. 2013. Highly expressed loci are vulnerable to misleading ChIP localization of multiple unrelated proteins. Proc Natl Acad Sci USA 110: 18602–18607.

Thomas-Chollier M, Darbo E, Herrmann C, Defrance M, Thieffry D, van Helden J. 2012. A complete workflow for the analysis of full-size ChIP-seq (and similar) data sets using peak-motifs. Nat Protoc 7: 1551–1568.

Tsankov AM, Gu H, Akopian V, Ziller MJ, Donaghey J, Amit I, Gnirke A, Meissner A. 2015. Transcription factor binding dynamics during human ES cell differentiation. Nature 518: 344–349.

Vega VB, Cheung E, Palanisamy N, Sung W-K. 2009. Inherent signals in sequencing-based Chromatin-ImmunoPrecipitation control libraries. PLoS ONE 4: e5241.

Wang J, Jia ST, Jia S. 2016. New Insights into the Regulation of Heterochromatin. Trends Genet 32: 284–294.

Xing H, Mo Y, Liao W, Zhang MQ. 2012. Genome-wide localization of protein-DNA binding and histone modification by a Bayesian change-point method with ChIP-seq data. PLoS Comput Biol 8: e1002613.

Xu H, Handoko L, Wei X, Ye C, Sheng J, Wei C-L, Lin F, Sung W-K. 2010. A signal-noise model for significance analysis of ChIP-seq with negative control. Bioinformatics 26: 1199–1204.

Xu H, Wei C-L, Lin F, Sung W-K. 2008. An HMM approach to genome-wide identification of differential histone modification sites from ChIP-seq data. Bioinformatics 24: 2344–2349.

Zhang Y, Liu T, Meyer C, Eeckhoute J, Johnson D, Bernstein B, Nusbaum C, Myers R, Brown M, Li W, et al. 2008. Model-based Analysis of ChIP-Seq (MACS). Genome Biol 9: R137.

